# Gene exchange networks define species-like units in marine prokaryotes

**DOI:** 10.1101/2020.09.10.291518

**Authors:** R. Stepanauskas, J.M. Brown, U. Mai, O. Bezuidt, M. Pachiadaki, J. Brown, S.J. Biller, P.M. Berube, N.R. Record, S. Mirarab

## Abstract

Post-submission note. Since the original submission of this manuscript to bioRxiv, we discovered that some of our results may be impacted by the limitations of some of the comparative genomics tools used in this study. We are working on a revised version of the manuscript.

Although horizontal gene transfer is recognized as a major evolutionary process in Bacteria and Archaea, its general patterns remain elusive, due to difficulties tracking genes at relevant resolution and scale within complex microbiomes. To circumvent these challenges, we analyzed a randomized sample of >12,000 genomes of individual cells of Bacteria and Archaea in the tropical and subtropical ocean - a well-mixed, global environment. We found that marine microorganisms form gene exchange networks (GENs) within which transfers of both flexible and core genes are frequent, including the rRNA operon that is commonly used as a conservative taxonomic marker. The data revealed efficient gene exchange among genomes with <28% nucleotide difference, indicating that GENs are much broader lineages than the nominal microbial species, which are currently delineated at 4-6% nucleotide difference. The 42 largest GENs accounted for 90% of cells in the tropical ocean microbiome. Frequent gene exchange within GENs helps explain how marine microorganisms maintain millions of rare genes and adapt to a dynamic environment despite extreme genome streamlining of their individual cells. Our study suggests that sharing of pangenomes through horizontal gene transfer is a defining feature of fundamental evolutionary units in marine planktonic microorganisms and, potentially, other microbiomes.

## MAIN TEXT

Horizontal gene transfer (HGT) is a key evolutionary process in *Bacteria* and *Archaea*, with impacts ranging from the spread of antibiotic resistance in human pathogens to the emergence of new players in planetary biogeochemical cycles ^1-5^. While many HGT studies have focused on remarkable yet rare gene transfers between evolutionarily distant microorganisms, routine HGT among close relatives has also been reported in the ocean ^6-9^ and other environments ^10-14^. The majority of microbial genes are flexible, i.e. they are not present in all members of a given lineage, which suggests frequent exchange ^15,16^. However, the identification of HGT phylogenetic boundaries and predominant sources, as well as the estimation of HGT rates in nature have proven difficult ^1-5,10,11^. Most flexible genes are rare ^17,18^, thus studies of their distribution in specific organisms may require bias-free genome sampling at scales that remain hard to achieve. Furthermore, homologous recombination, one of the key steps in HGT, is the most frequent among near-identical genomes ^10,19^, where it is also the hardest to detect using computational tools that rely on DNA sequence discrepancies ^12,20^. Yet another challenge is complex and poorly understood microbial biogeography, which acts from microscopic to global scales and obscures the discrimination of the effects of evolutionary distance and encounter rates on the distribution of genetic material in a population ^21,22^. Consequently, ecosystem-wide and global patterns of HGT remain largely unknown. Likewise, although many theoretical concepts of microbial evolution view HGT as essential to speciation ^21^, such concepts have not replaced arbitrary species demarcations in contemporary microbial taxonomy ^23-25^, due to limited empirical support.

Marine planktonic *Bacteria* and *Archaea* (prokaryoplankton) play essential roles in the global carbon cycle, nutrient remineralization and climate formation and constitute two thirds of ocean’s biomass ^26-29^. Prokaryoplankton inhabiting epipelagic environments in tropical and subtropical latitudes are efficiently dispersed around the globe by ocean currents and differ genomically from prokaryoplankton inhabiting higher latitudes and greater depths ^17,30^. Thus, they form a global microbiome with clear geographic boundaries that has been evolving over geological time scales. Here, we analyze 12,715 randomly sampled single amplified genomes (SAGs), called GORG-Tropics, recovered from 28 globally distributed samples of tropical and subtropical, euphotic ocean water ^17^. Initial analyses found, remarkably, that no two of these cells had identical gene repertoire ^17^, indicating that HGT may play a major role in shaping these genomes. We leverage the large size and randomized genome sampling approach of GORG-Tropics to analyze how HGT relates to microbial evolutionary distance in this global, well-mixed microbiome.

### Flexible genes

The distribution of flexible genes is expected to be randomized in a population undergoing frequent HGT ^15,16^. In search for genealogical boundaries of HGT, we analyzed how differences in gene content relate to genome nucleotide difference (GND) – a commonly used metric of evolutionary distance calculated as 1-ANI (average nucleotide identity) ^10,19,31,32^ – in pairs of GORG-Tropics genomes. Surprisingly, we observed a non-linear relationship, with a distinct breakpoint at 27.6% GND **(Fig. 1A)**. Below this threshold, the fraction of non-orthologous genes increased only slightly with increasing GND, while a strong, positive relationship emerged at evolutionary distances above this GND value. The saturation of codon wobble positions could not explain this discontinuity, because a similar, non-linear pattern emerged in the relationship between genome-wide amino acid difference (AAD) and the fraction of non-orthologous genes, with an inflection point of 39.3% AAD (**Fig. 1B)**. To confirm that these patterns were not an artifact of our computational tools, we searched for and found no discontinuity in the GND-AAD relationship at these values (**Fig. S1)**. Next, we generated a set of mock genomes with randomized gene content, which failed to produce discontinuities in the relationship between GND and fraction of non-orthologous genes (**Fig. S2A**). Furthermore, changing the amino acid identity threshold in the CompareM ^33^ ortholog identification algorithm from 30% (default) to 50% had only a minor impact on the breakpoint in GORG-Tropics (**Fig. S2A**). Finally, we replaced the CompareM *de novo* ortholog search with identification of shared hidden Markov model-based Prokka ^34^ annotations, which produced a similar pattern with a near-identical breakpoint (**Fig. S2B**). This indicates that discontinuities at ∼28% GND and ∼39% AAD are intrinsic features of prokaryoplankton in the tropical and subtropical ocean. Below these thresholds, GND has little influence on the fraction of non-orthologous genes, which averaged at 29% and was similar to the average of 28% of strain-specific genes found in well-characterized, nominal bacterial species ^35^. Similar discontinuities, although at slightly higher GND values, were observed in pairwise differences in genome size and G+C content (**Fig. 1E**). We propose that the most plausible explanation for the observed discontinuity is HGT frequency being sufficient to maintain shared pools of flexible genes (pangenomes) among bacterioplankton cells below ∼28% GND. Above this approximate threshold, insufficient frequency of HGT fixation leads to the separation of pangenomes and accumulation of gene content differences in individual cells.

**Fig. 1.**
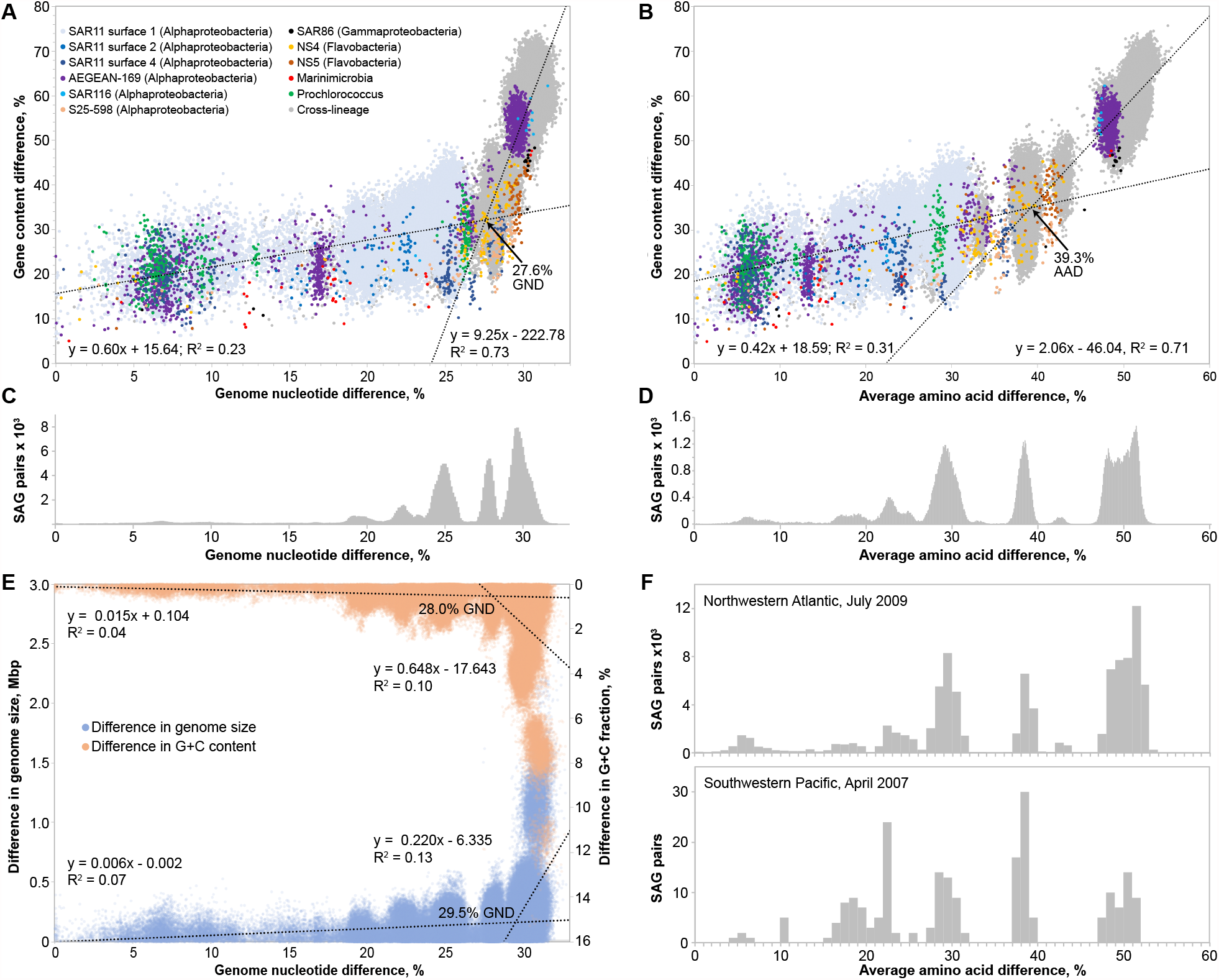
Patterns of genome divergence of marine bacterioplankton. Fraction of non-orthologous genes in SAG pairs in relation to their genome nucleotide difference (GND) (A) and amino acid difference (AAD) (B). Frequency distribution of GND (C) and AAD (D), calculated at 0.1% GND and AAD intervals. (E) Differences in estimated genome size and G+C content in SAG pairs in relation to GND. (F) Frequency distribution of AAD in SAGs from two locations and sample collection times, calculated at 1% AAD intervals. To minimize analytical uncertainties, we sub-sampled GORG-Tropics to 819 bacterial genomes with ≥80% estimated genome completion that contain 16S rRNA genes. In panels A and B, colors of data points designate taxonomic lineages represented by ≥30 genome pairs. Orthologs were identified using a 30% amino acid identity threshold with CompareM ^33^. In panel F, Northwestern Atlantic is represented by field samples SWC-09 and SWC-10, while Southwestern Pacific is represented by field samples 117-Fiji-Hawaii-2007, 121-Fiji-Hawaii-2007, 133-Fiji-Hawaii-2007 and 137-Fiji-Hawaii-2007 ^17^. Regression models and points of inflection were estimated by change point regression and segmented models ^51^.

The distribution of GORG-Tropics SAG pairs along the GND and AAD gradients exhibited multiple sharp peaks averaging ∼ 19.5%, 22.5%, 25%, 28% and 30% GND (**Fig. 1 C-D**). These peaks did not coincide with breakpoints in relationships between GND, AAD and gene content (**Fig. 1 A-B**), indicating that mechanisms causing these peaks had no major impact on our estimated HGT boundaries. The low frequency of genome pairs with near-0% GND suggests that GND distribution peaks were not caused by multiple lineages forming blooms of near-clonal populations during the GORG-Tropics sample collection. Based on the finding of the same peaks in geographically and temporally distant samples (**Fig. 1F**) and their mixed taxonomic composition (**Fig. 1A-B**), we speculate that these peaks reflect concurrent population expansions of multiple lineages during ancient, ecosystem-wide perturbations, and deserve further investigation. As reported previously ^17^, fewer than 0.1% of GORG-Tropics SAG pairs fell below the 4% GND threshold for the contemporary, nominal definition of microbial species ^24^. This contrasts with several recent reports of elevated abundance of microorganisms with near-0% GND, which were interpreted as empirical support of the contemporary species delineations ^36,37^. Although differences between marine and non-marine microbiomes may play a role, this discrepancy among studies also underlines the potential importance of genome selection strategies: a) randomized sample of individual cells ^17^, b) public databases dominated by specific taxa ^36^, and c) metagenome bins defined by assembly and binning algorithms ^37^.

### Core genes

Frequent HGT of core genes (genes that are present in all members of a lineage) is expected to randomize the distribution of their alleles in a population. We utilized GORG-Tropics to study how GND relates to the divergence of two highly conserved, universal genes that are broadly utilized in phylogenetic analyses: 16S rRNA ^38^ and *rpsC* ^39^. In various bacterioplankton lineages, divergences of both genes correlated with GND only weakly at low GND values, indicating frequent HGT **(Fig. 2 A-B)**. We found completely identical 16S rRNA nucleotides and predicted RpsC proteins in cell pairs with up to 14% and 19% GND. In contrast, more evolutionarily distant SAG pairs produced strong correlations of GND versus 16S rRNA and RpsC divergences, with inflection points of these non-linear relationships approximating 23% and 22% GND. Uncorrelated variable rates of evolution among lineages could not explain this pattern, because an opposite trend (loss of correlation with increasing GND) was observed in a simulation with randomly varied evolution rates **(Fig. S3)**. This suggests that HGT frequency of 16S rRNA and *rpsC* genes is sufficient to counteract the accumulation of nucleotide substitutions among cells that fall below these approximate divergence thresholds. Further support for this conclusion is provided by the high levels of discordance between 16S and 23S rRNA gene phylogenies at low GND and low levels of discordance at high GND, with the breakpoint at ∼22% GND (**Fig. 2C-D**). This implies frequent recombination of rRNA operon segments among cells with <22% GND.

**Fig. 2.**
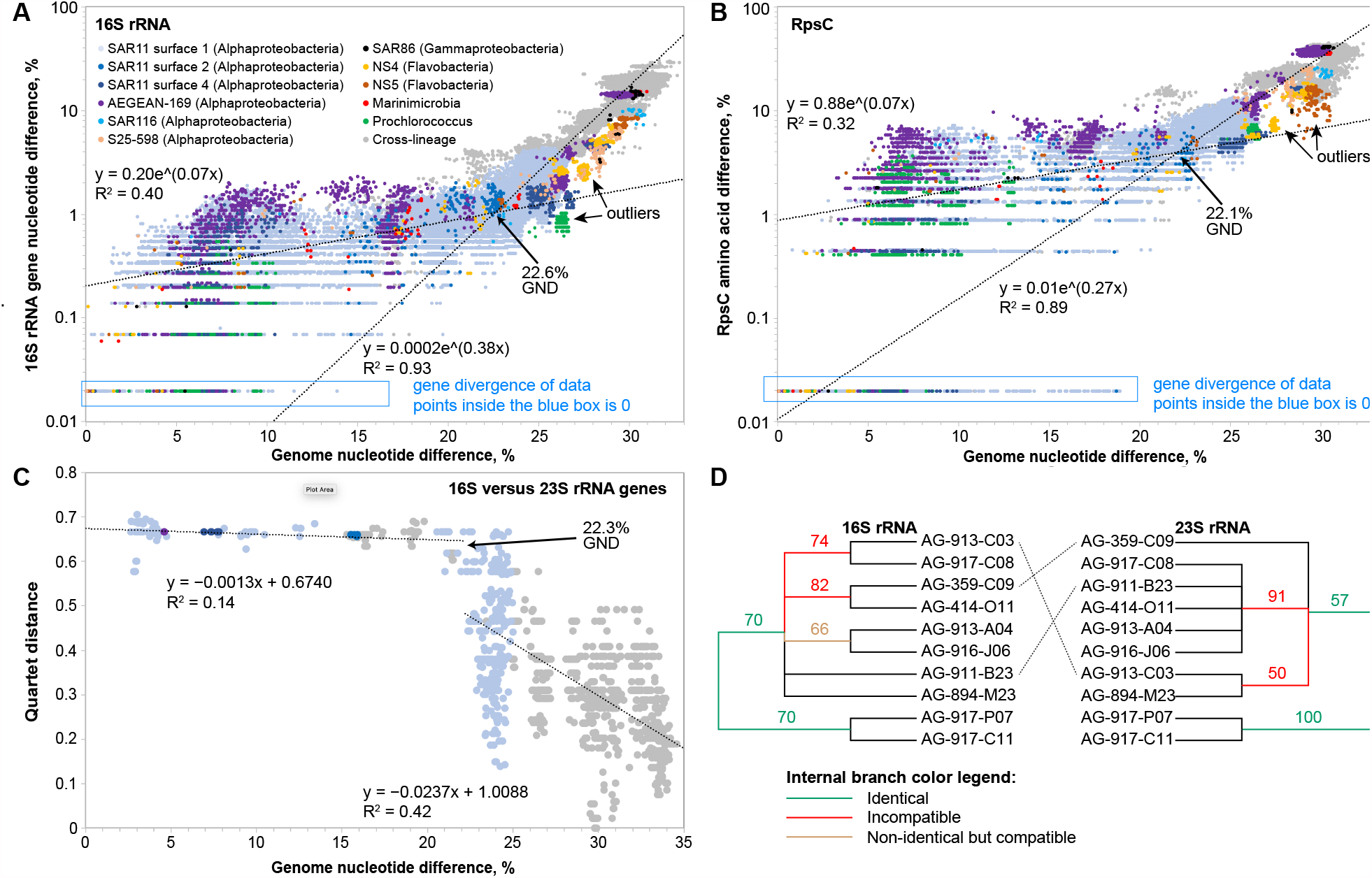
Divergence of selected core genes in relation to genome divergence. Relationships between genome nucleotide difference (GND) and sequence divergences of 16S rRNA (A) and RpsC (B) genes. (C) Phylogenetic discordance of 16S and 23S rRNA gene trees along GND gradient. Each dot represents a subset of 10 SAGs, characterized by its mean pairwise GND. Tree discordance is measured using normalized quartet distance after collapsing branches with <70% bootstrap support. (D) Example of discordance of 16S and 23S rRNA gene phylogenies among SAR11 surface-1 lineage SAGs with mean 2.9% GND. Numbers by nodes indicate bootstrap support. Branches with <50% bootstrap support were removed. All genome labels are aligned except those indicated by dotted lines. Regression models and points of inflection were estimated by change point regression ^51^ using segmented (A and B) and stegmented (C) models. Included are 819 GORG-Tropics SAGs with ≥80% estimated genome completion and the recovery of the 16S rRNA gene. In panels A and B, colors of data points designate taxonomic lineages represented by ≥30 genome pairs.

Interestingly, a discernible subset of phylogenetically diverse genome pairs formed correlation patterns that ran parallel to the bulk of data points, with unusually low 16S rRNA and RpsC divergences relative to their GND (marked as “outliers” in **Fig. 2 A-B**). We hypothesize that such outliers form as a result of divergent GND thresholds for the frequent HGT of highly conserved genes (**Fig. 2**) versus genome average (**Fig. 1**), leading to differences in the evolutionary breadth of populations that share pangenomes of various genes.

The rRNA operon is generally considered one of the most stable and slowest-evolving genome elements and therefore has served as the foundation for microbial genealogy inferences since the 1970s ^40^. This assumption was challenged by several studies that detected HGT of rRNA genes in closely related strains and produced functional ribosomes from mosaic rRNA *in vitro* ^12^. Our results help reconcile these conflicting findings by identifying the specific GND range in which the fixation of horizontally transferred rRNA genes may be frequent in marine prokaryoplankton. The results of GND comparisons with 16S rRNA and *rpsC* divergences indicate that even core genes are involved in frequent HGT among close relatives. This conclusion corroborates with previous reports of recombination of core genes is SAR11, the most abundant lineage of marine prokaryoplankton ^6^. We hypothesize that the consistent presence of core genes in closely related microorganisms is not caused by the exclusion of these genes from HGT, as often assumed ^33^, but rather by a dynamic process that involves HGT and selection against cells without compatible variants of these genes.

### Recent HGT

We counted protein-coding genes with 0%, ≤1%, and ≤5% nucleotide difference in genome pairs along the GND gradient as a proxy for putative, recent HGT events **(Fig. 3A)**. Observations of such genes declined exponentially from 0% to 27% GND range. The slope of the exponential decline was similar among the various COG categories, although the breakpoint varied, with categories P (inorganic ion transport and metabolism) and F (nucleotide metabolism and transport) dominating putative HGT counts at high GND **(Fig. 3B)**. Accordingly, prior studies have reported exponential declines of recombination rates among closely related bacterial ^10^ and archaeal ^19^ isolates along the GND gradient. Our findings of near-identical genes in genome pairs may be a result of both HGT and variable rates of nucleotide substitution among genes, as the two factors are difficult to disentangle. Nevertheless, it is interesting that the GORG-Tropics dataset, which contains a wide spectrum of microbial diversity, exhibited a non-linear decline in near-identical gene counts at GND > 28%. This discontinuity was observed at the same GND value when using either 0%, 1% or 5% gene divergence thresholds, indicating a robust, biological feature rather than a methodological artifact. The accelerated decline in counts of near-identical genes at 28% GND coincided with the discontinuity in gene content similarity at the same GND value (**Fig. 1**), in support of infrequent HGT among cells with >28% GND.

**Fig. 3.**
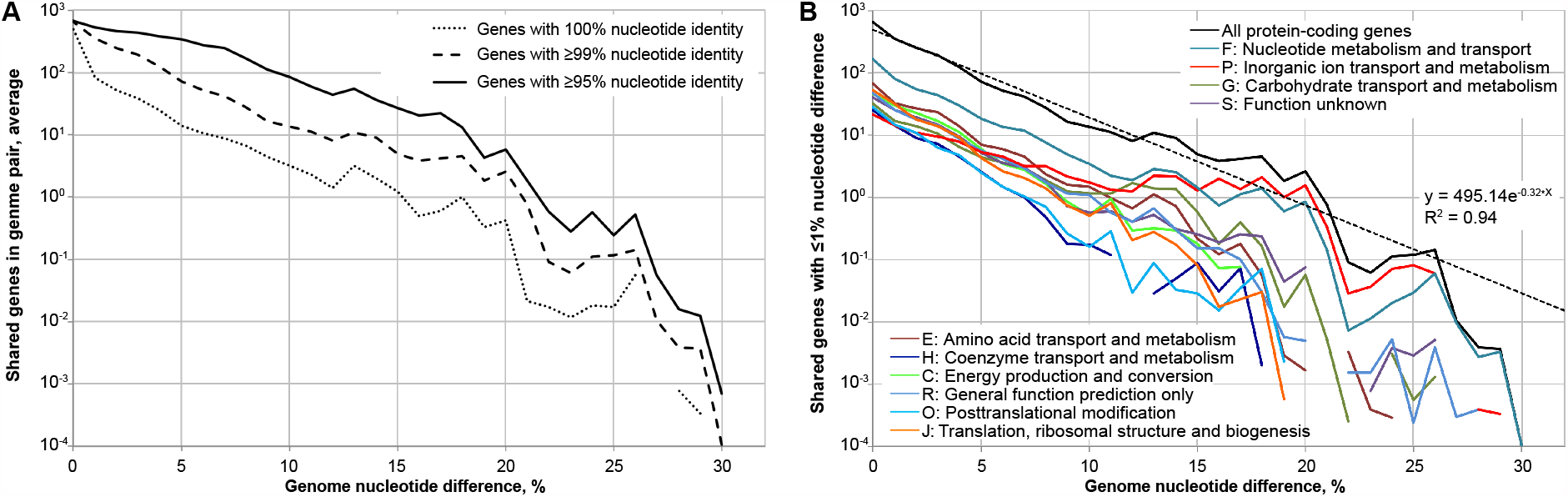
Average counts of genes with high nucleotide identity in pairs of GORG-Tropics SAGs as a function of GND. Included are 819 GORG-Tropics SAGs with ≥80% estimated genome completion and the recovery of the 16S rRNA genes. Counted are only gene alignments that cover ≥80% of query sequence length. **(A)** Average counts of protein-coding genes with 95-100% nucleotide identities in genome pairs. **(B)** Average counts of protein-coding genes with ≥99% nucleotide identity in genome pairs. Colored datasets correspond to 10 most abundant COGs. Regression is based on the 0-27% GND range. Data discontinuities correspond to values equal to zero.

### Gene exchange networks

Using the 28% GND threshold, we clustered GORG-Tropics SAGs into groups that we term gene exchange networks (GENs) according to single-linkage clustering. These GENs vary in the extent of internal linkage, ranging from 11% to 100% (mean 78%). Of the 12,715 GORG-Tropics SAGs, 9,176 SAGs formed 168 GENs of ≥2 members (**Figs. 4, S4**). Of the 3,539 SAGs remaining as singletons, 94% had assembly sizes <0.5 Mbp (**Fig. S5**), suggesting that their exclusion from GENs is mostly due to limited genome recovery. The 42 most populous GENs accounted for >90% of the 7,208 SAGs with ≥0.5 Mbp assemblies. Given the good agreement in taxonomic and genomic compositions of GORG-Tropics SAGs and other marine prokaryoplankton ‘omics datasets ^17^, these 42 most populous GENs constitute a realistic representation of the majority of prokaryoplankton in the tropical and subtropical ocean.

**Fig. 4.**
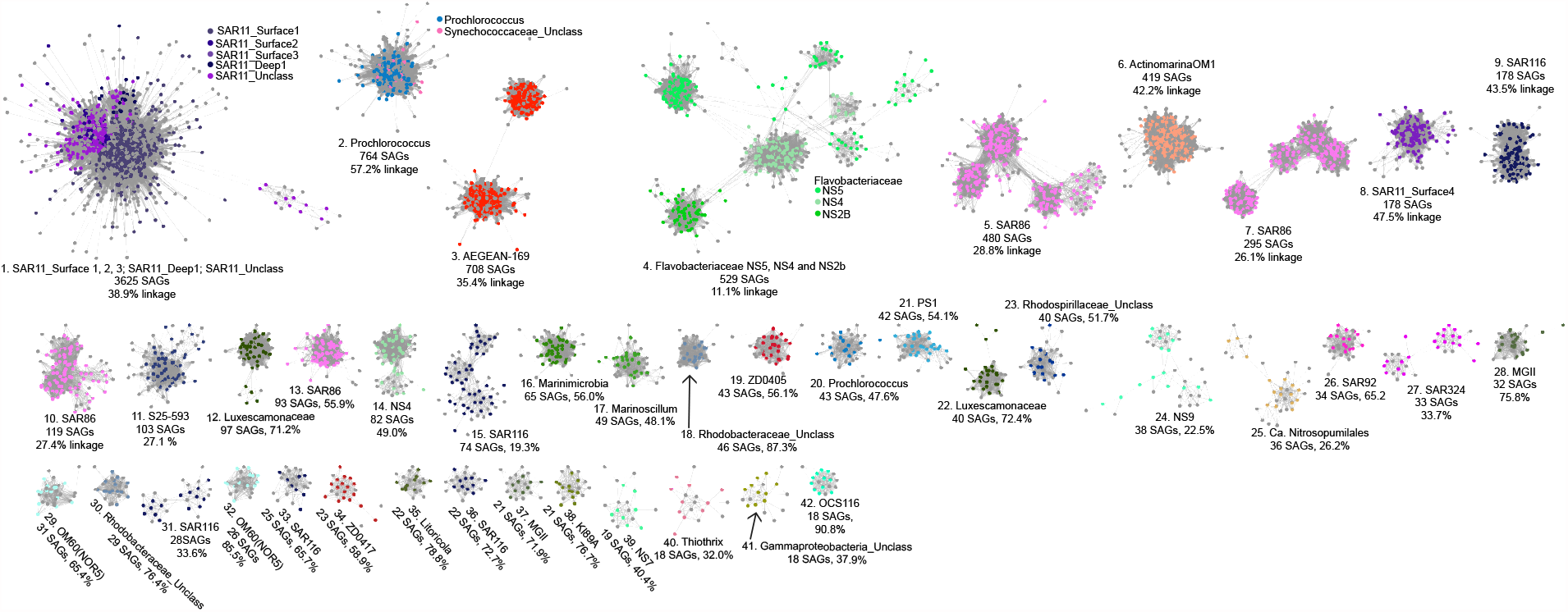
The 42 GENs that account for 90% of GORG-Tropics SAGs. Nodes represent SAGs, edges represent GND ≤ 28%, node colors and titles represent literature-derived phylogenetic-ecological lineages. Text indicates GEN ID, 16S rRNA-based lineage, count of member SAGs and network linkage (ratio of actual versus possible link counts). Additional information on a complete set of GENs can be found in Fig. S4 and Data S1 and S2.

Most GENs contained only one of the previously described lineages of marine prokaryoplankton (**Figs. 4, S4**), demonstrating good agreement between GENs and the earlier, primarily 16S rRNA-based phylogenetic demarcations, with a few notable exceptions. The largest GEN, comprised of 3,625 SAGs, encompassed all SAGs of SAR11 (Alphaproteobacteria) ecotypes surface-1, surface-2, surface-3, deep-1 and “unclassified”, indicating that these lineages share a single pangenome, although with some internal sub-clustering (**Fig. S4**). This observation helps explain an earlier finding of rhodopsin, a light-harvesting system, being encoded in ecotype deep-1, which is found primarily below the euphotic depths ^41^. Meanwhile, SAR11 surface-4 formed a separate GEN, which agrees with the recent findings of divergent biosynthetic capabilities ^17^ and rRNA operon structure ^42^ of this lineage as compared to other SAR11. Our network analysis also indicated an incomplete separation of Flavobacteriaceae lineages NS2b, NS4 and NS5. By contrast, we found that many of the previously described lineages, which span genus to phylum taxonomic units, form multiple GENs. Notably, *Prochlorococcus* (Cyanobacteria) formed two large and several small GENs, indicating a separation of gene pools between the high-light adapted clades (e.g. HLI, HLII, HLIII, and HLIV) and the low-light adapted LLI clade. These data suggest that within the world’s most abundant lineage of phototrophs, there exist multiple GENs, each with likely separate evolutionary trajectories. The most abundant Gammaproteobacteria lineage SAR86 formed four GENs with >90 members, three of which had low linkage and substructures, indicating a high degree of genomic and evolutionary heterogeneity. Multiple GENs were also formed by lineages SAR116, PS1 (Alphaproteobacteria), OM60 (Gammaproteobacteria), SAR324 (Deltaproteobacteria), MGII (Crenarchaeota) and others. In general, the topology of GENs ranged from high linkage (e.g. the two largest GENs of *Prochlorococcus*) to a complex internal structure (e.g. the largest GENs of Flavobacteriaceae and SAR86), which is consistent with the gradual separation of sister GENs along the GND gradient (**Figs. 1, 2**). Thus, GENs offer a biologically meaningful and practical paradigm for studies of environmental microorganisms.

## Discussion

Our findings suggest that bacterioplankton in the tropical euphotic ocean form gene exchange networks that are characterized by shared pangenomes, with 28% GND as their approximate boundary (**Fig. 5**). Extensive HGT within GENs helps explain how the ∼40 million gene types, most of which rare ^17,18^, are maintained by bacterioplankton lineages that are characterized by extreme genome streamlining ^43^, with an average genome size of 1.6 Mbp ^17^. Access to a large pool of readily exchanged genes within GENs likely facilitates adaptations of marine bacterioplankton to a dynamic environment despite the limited coding potential of their individual cells.

**Fig. 5.**
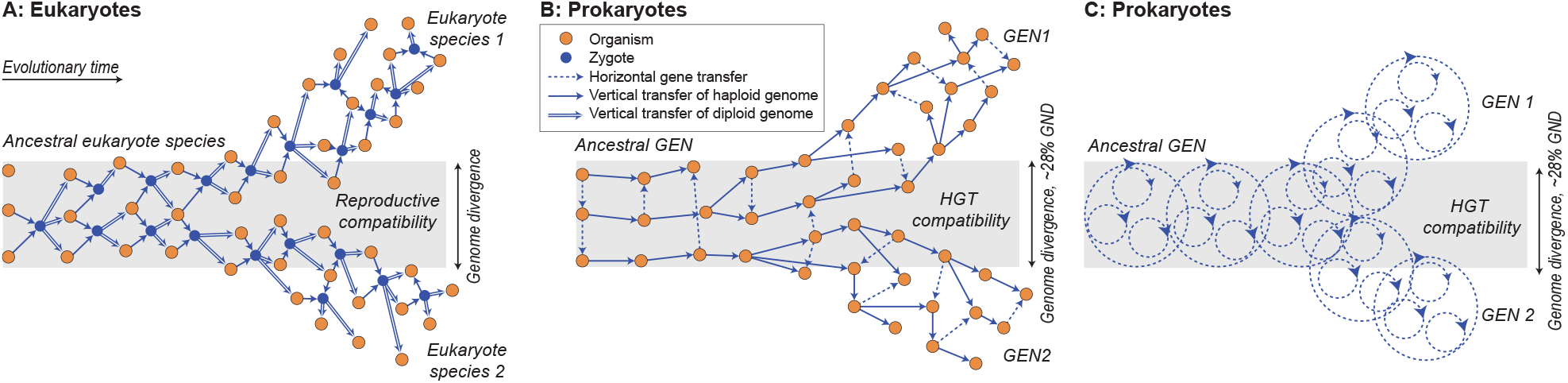
Conceptualized gene flow and divergence in sexually reproducing eukaryotes (A) and prokaryoplankton GENs (B, C). In contrast to the formation of a zygote in eukaryotes, prokaryote HGT is not coupled to reproduction and is unidirectional. Furthermore, since HGT involves both homologous and non-homologous recombination, individual members of GENs contain only a fraction of GEN’s pangenome. A and B depict gene flow among cells. C depicts a broader, nested/fractal pattern of gene flow within GENs.

The observed GEN boundaries are soft, indicating that the transition from HGT-driven GEN cohesion to pangenome separation between sister GENs takes place over considerable evolutionary time (**Fig. 1**) and is not uniform across gene types (**Figs. 2, 4**). Therefore, a fixed threshold in GEN delineation would not be an accurate reflection of natural evolutionary processes. Nevertheless, the soft breakpoint that averages ∼28% GND demonstrates that marine bacterioplankton GENs exceed the evolutionary breadth of current, nominal definitions of bacterial species, which range between 4-6% GND and are designed to maintain the continuity of historical taxonomies ^23-25^. Assuming an average amino acid substitution rate of ∼0.05% per Myr ^44^, the age of bacterioplankton GENs likely ranges in hundreds of Myr. This time scale is consistent with the estimated age of *Prochlorococcus*, which forms two of the largest bacterioplankton GENs, and human commensal lineages for which reliable calibration of the molecular clock is available ^44,45^. The longevity of GENs helps explain the efficient, global exchange of genes within marine GENs, because the entire ocean water volume is overturned by the global thermohaline circulation every 1-2 kyr, while surface ocean currents operate at substantially shorter timeframes ^46^.

Although GORG-Tropics did not capture putative HGT events among cells with >32% GND, this does not mean that such events never occur, as exemplified by prior reports of cross-domain HGT ^47^. Some gene flow among evolutionarily distant lineages may also take place through intermediary lineages. However, the rate of fixation of genes acquired across GEN boundaries may be insufficient to obscure the predominant evolutionary patterns, akin to the rare introgression events in eukarya ^48^. Furthermore, some ancient HGT between now evolutionarily distant lineages may have occurred while both the donor and the recipient were still part of the same GEN. Likewise, multiple studies have demonstrated distinct patterns of HGT within substantially more narrow lineages of marine microorganisms, variably named “ecotypes”, “genotypes”, “backbones” and “populations”^8,9,49^. Our data suggest that that such lineages may not maintain isolated pangenomes, therefore their divergence from sister lineages may be temporary and reversible. We speculate that HGT manifests at multiple levels, with impacts of evolutionary proximity, encounter rates and HGT mechanisms producing nested, fractal-like patterns of gene flow in complex microbiomes **(Fig. 5C)**.

Efficient, global mixing of the euphotic, tropical ocean simplifies studies of microbiome-wide HGT in this environment. Dispersal and cell encounter limitations likely have a more profound impact on HGT patterns in environments such as soils and organismal microbiomes. However, the lifespan of GENs ranging in hundreds of millions of years may enable sufficient HGT to maintain global GENs even in such compartmentalized microbiome types. Accordingly, discontinuities in genome content along nucleotide divergence gradients are apparent in several earlier reports analyzing diverse microbial genome collections, although evolutionary origins of these discontinuities were not discussed ^31,50^. Our findings suggest that sharing of pangenomes through HGT defines fundamental evolutionary units in marine prokaryoplankton and demonstrates a novel approach to explore general HGT patterns in other microbiomes. Viewed as cohorts of cells with dynamic, shared gene pools, GENs offer a new conceptual framework to guide studies of microbial evolution, ecology and biotechnological exploration.

## SUPPLEMENTAL INFORMATION

Materials and Methods

Figures S1-S6

Tables S1-S2

## ACKNOWLEDGMENTS

We thank Elizabeth Fergusson, Brian Thompson, Corianna Mascena and Ben Tupper at the Bigelow Laboratory for Ocean Sciences’ Single Cell Genomics Center for the generation of single cell genomic data. We also thank Daniel Buckley (Cornell University) and Thane Papke (University of Connecticut) for valuable advice.

## AUTHOR CONTRIBUTIONS

R.S. developed the concept, managed the project and led manuscript preparation. J.M.B. performed most data analyses related to key genome characteristics and clustering. U.M. and S.M. performed phylogenetic discordance analyses and variable evolutionary rate modeling. O.B. performed putative HGT identification. N.R.R. performed change point regression analyses. M.P. and J.B. led data management and curation. P.M.B. and S.J.B. oversaw field sample collection and selection. All authors contributed to data interpretation and manuscript preparation.

## MATERIALS AND METHODS

### Data source

The main dataset utilized in this study is Global Oceans Reference Genomes Tropics (GORG-Tropics), which consists of 12,715 partial genomes of individual cells of bacteria and archaea obtained through a random selection from 28 globally distributed samples of tropical and subtropical, photic ocean water column ^1^.

### Genome-wide comparisons

Average nucleotide identity (ANI) of genome pairs was calculated using the ANIb method within pyani ^2^. Although computationally more demanding than alternative approaches, this technique offers superior sensitivity in a broader range of GND values ^3^. Due to high computational demands of this software, we limited this analysis to those 819 GORG-Tropics SAG assemblies that contained 16S rRNA gene sequences and were estimated to have ≥80% completion. ANI comparisons with >3% overlap in both directions were kept for analysis, self-comparisons were removed. Genome nucleotide difference (GND) was calculated as GND = 1-ANI. Detection of orthologous genes and amino acid identity were determined with CompareM, which uses reciprocal best hits to identify orthologous pairs ^4^. Search for nearly identical genes was done using reciprocal BLASTn ^5^ between the open reading frames of genome pairs. Mock genomes were generated by randomly sampling 10kb fragments from 16S containing SAGs that were ≥ 80% complete, to construct a set of 1,000 genomes that followed genome length distributions similar to the real data. Inflection points and model fits were estimated using change point regression using the segmented model, except for the regression analyses of phylogenetic quartet distance, where the “stegmented” model allowing for a jump between the two lines was used ^6^.

### Phylogenetic analyses

Multiple alignment of the 16S and 23S genes was performed using SINA ^7^ and followed by alignment trimming with trimAl version 1.4rev15 ^8^. Maximum likelihood trees were inferred using RAxML version 8.2.10 ^9^ with default settings and 1,000 bootstrap replicates. To compute discordance of 16S and 23S rRNA genes, we first created 3,000 genome subsets, each containing 10 SAGs. We used the single-linkage agglomerative clustering algorithm implemented in TreeN93 ^10^ to build a hierarchy of the genomes. At any threshold of GND, all genomes within that threshold belong to a sub tree of the single-linkage tree. We then cut the single tree at different thresholds of GND and chose 10 individual genomes at random from different subtrees. This allowed us to control the pairwise GND of the 10 genomes within a certain range. By varying the threshold between 0.5% and 29.5% (steps: 0.5), we obtained 60 groups of subtrees and randomly sampled 50 subsets of size 10 from each group. This procedure produced 3,000 subsets that covered the range between 2.5% and 34.6% mean pairwise GND (average: 18.9%). Once the subsets of genomes were selected, we restricted the original 16S and 23S trees to include only the 10 selected SAGs. We computed the bootstrap support for the restricted trees by restricting the bootstrap replicate trees to the 10 selected SAGS and computing bipartition support. We then collapsed all branches with BS<75%. We discarded all pairs of subtrees that had more than 2/3 quartet trees (140 out of 210 possible) as unresolved. This left us with 1,422 pairs of subtrees. For each pair, we computed the normalized weighted quartet distance. A quartet match score was computed where each quartet was counted as 1 if resolved identically in both trees, as 0 if resolved differently in the two trees, and 1/3 if unresolved in one or both trees (because there is a 1/3 chance for trees to resolve the quartet identically at random). The weighted quartet distance was computed by subtracting the total match score from the total quartets and then normalizing by the total number of quartets. To compute quartet matches, we used tqDist^11^.

To test if a simple model of rate variation can describe patterns of GND discordance that we observed, we used a simulation. We first inferred a concatenation tree by utilizing the GToTree phylogenomic workflow ^12^ using an HMM set of 74 single copy gene targets. A Maximum likelihood tree was generated using FastTree version 2.1.10 ^13^ with the default parameters. We then rooted the tree using MV rooting ^14^ and made the tree ultrametric (i.e., clock-like) using wLogDate ^15^, fixing the tree height to 1. We then multiplied each branch length by a rate multiplier drawn randomly from the Gamma distribution with shape and scale set to 1 (thus, mean: 1; variance: 1). This model is used in many existing simulation methods for capturing rate variation across genes ^16,17^. The rate multipliers represent changes in evolutionary rate and are modelled here as being independent. We repeated this process 5 times, obtaining 5 potential “gene trees” that differ from the main SAG tree only in terms of rates of evolutionary change. We then compared all pairwise distances in all pairs of these simulated gene trees and used these values to draw **Fig. S3**.

### Network analysis

To construct GND-based network clusters for GORG-Tropics, we first used the results from the pyani ANIb comparison ^2^ computed for GORG-Tropics SAG assemblies that had ≥80% completion estimates and contained 16S rRNA gene sequences. This provided us with GND estimates ranging ∼0-32%. In order to complement these results with additional SAGs, we took advantage of the computational efficiency of fastANI ^3^, which we ran on all GORG-Tropics to estimate GND up to ∼ 25% between SAG pairs. ANI estimates for the two calculators was linear for values below 20% GND (R^2^ = 0.992, fastani = ani_b * 0.977 + 2.4). Above 20% GND, fastani values leveled out at GND = 25%, while ANIb values calculated by pyani extended above 25% GND. The pyani and FastANI results were combined and networks were identified as single-linkage network graphs. Sub-modules were identified using the “community” network clustering algorithm within the Louvain Python package ^18^ and visualized with cytoscape ^19^.

## SUPPLEMENTAL INFORMATION

**Fig. S1.**
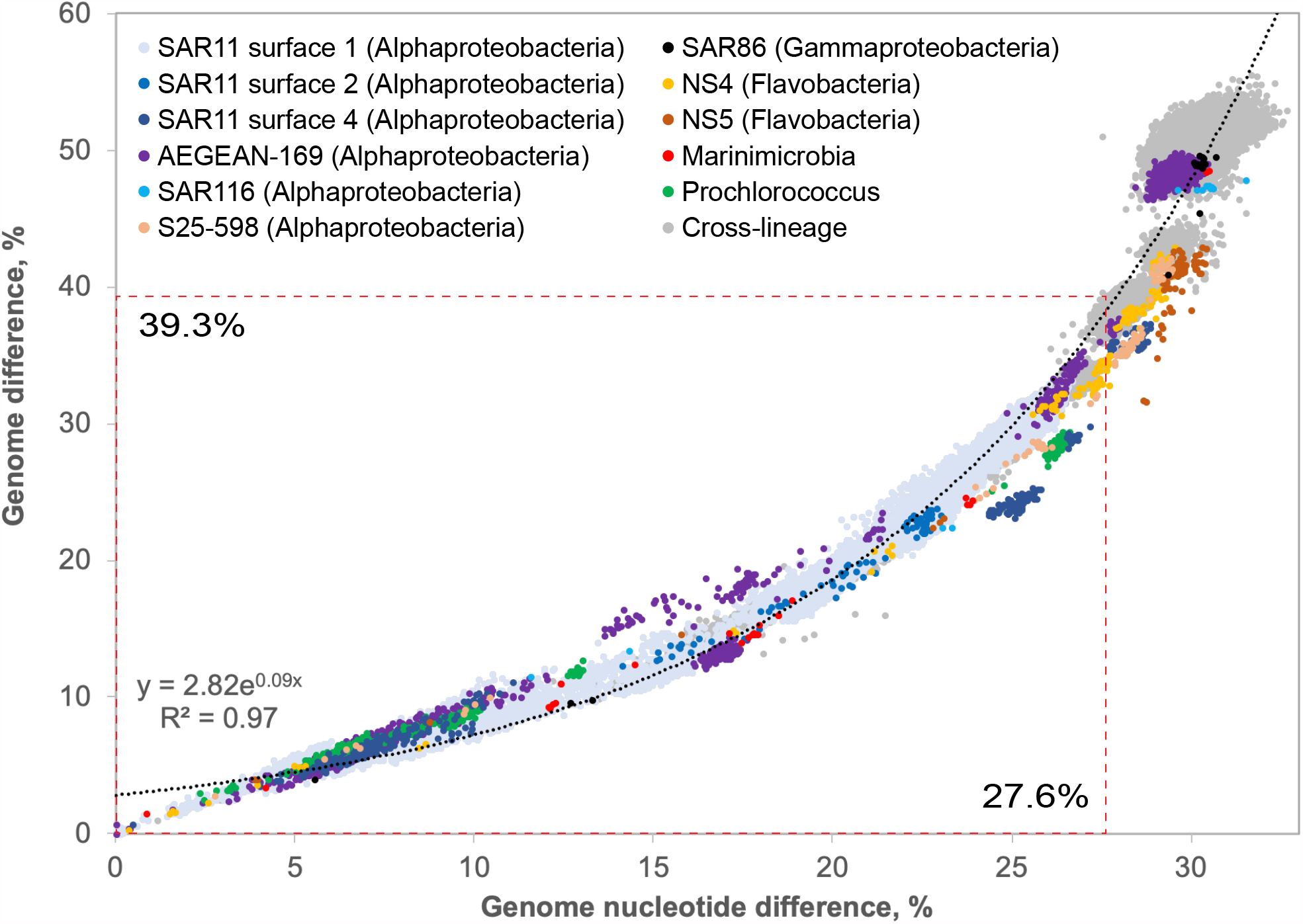
Relationship between GND and AAD. Red-dotted lines indicate inflection points observed in Fig. 1.

**Fig. S2.**
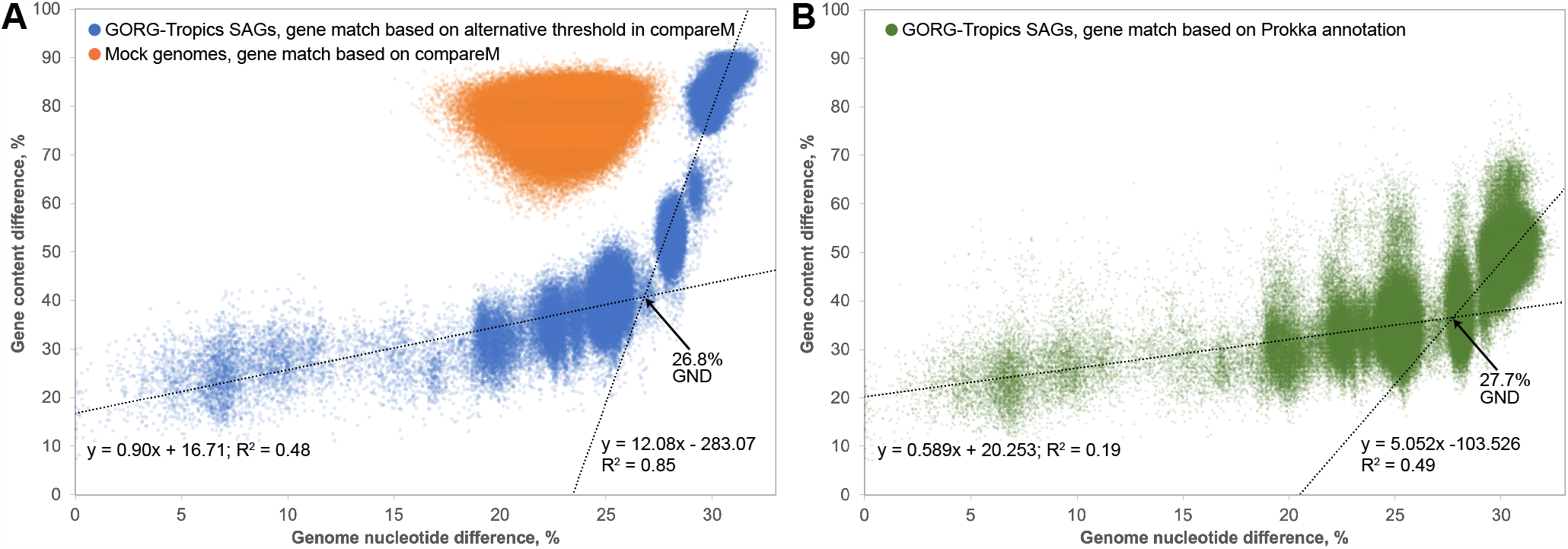
Relationship between GND and the difference in protein-coding gene content. (A) CompareM-based gene content differences: mock genomes analyzed using default settings (orange) and GORG-Tropics SAGs analyzed using a 50% amino acid identity threshold in ortholog detection. (B) GORG-Tropics SAGs analyzed with Prokka.

**Fig. S3.**
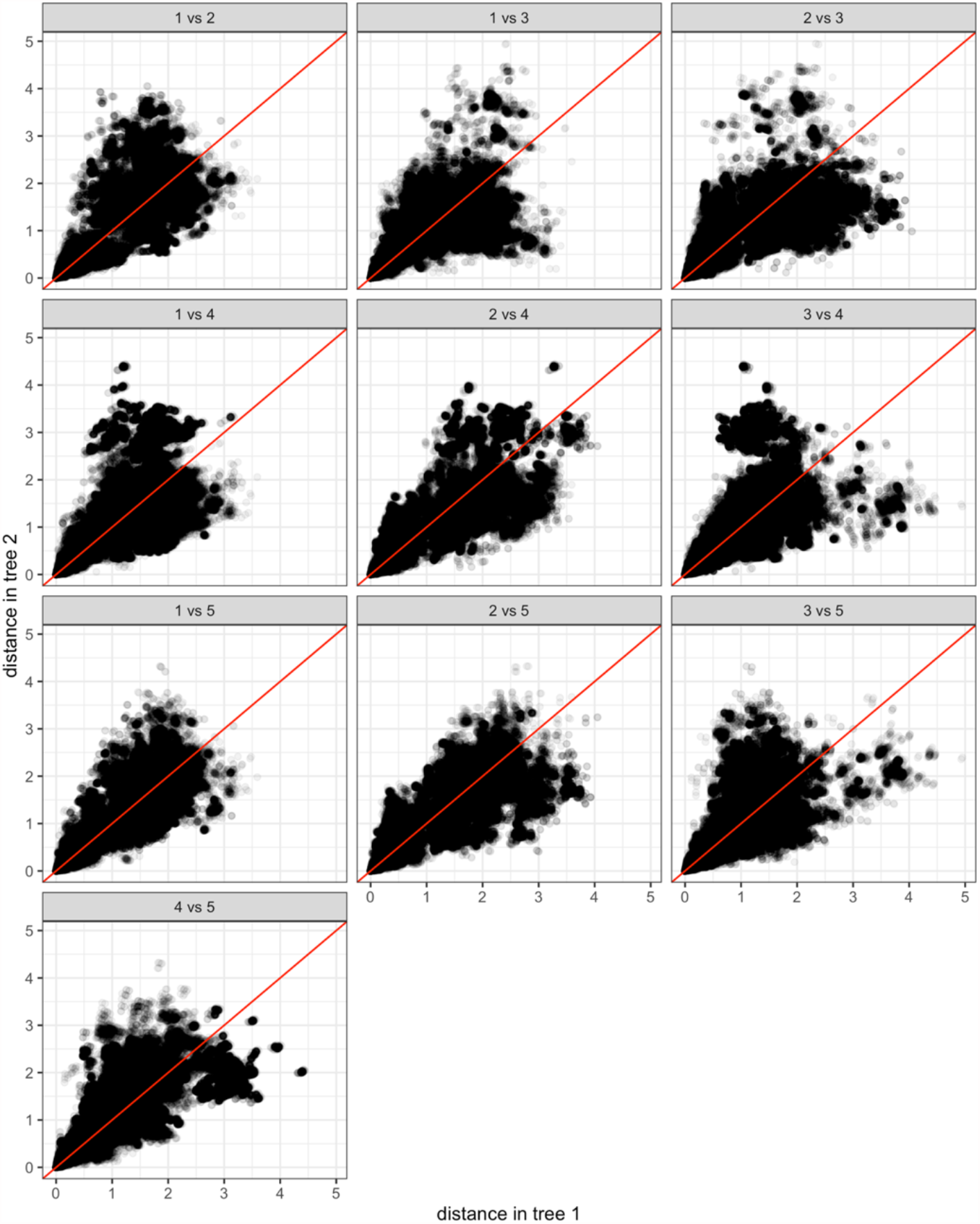
Simulation of varied evolution rates. We show the correlation between branch lengths between two simulated gene trees, showing the total tree distance for all pairs of leaves, each represented as one dot. The gene trees are simulated by starting from a common tree, keeping the topology fixed, but simply multiplying all branch lengths by a rate multiplier drawn randomly from the Gamma distribution (shape and scale both set to 1). This model is used in many existing methods for capturing rate variation across genes and is found to be a good match to empirical data ^1,2^.

**Fig. S4.**
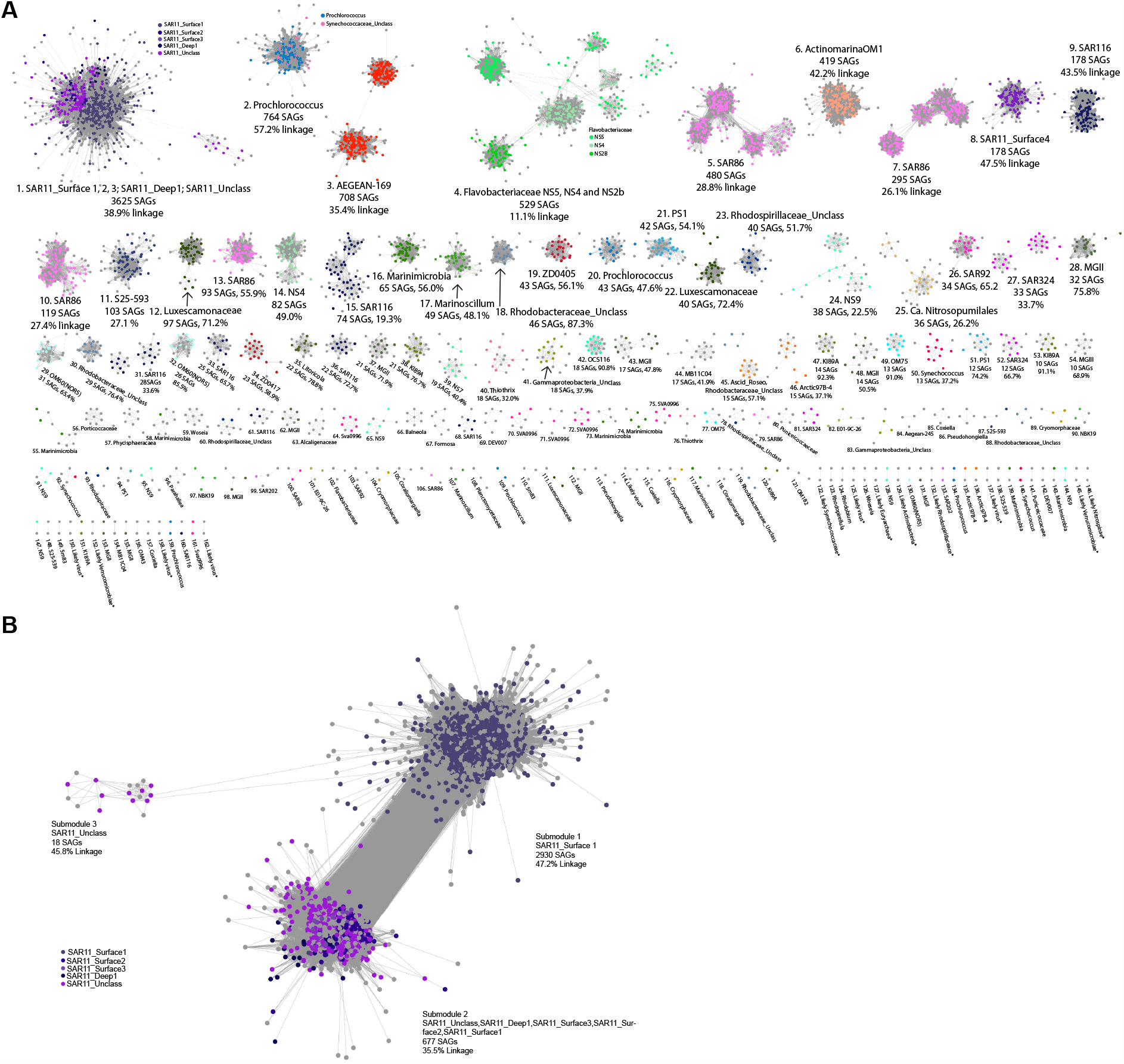
Gene exchange networks (GENs). Nodes represent SAGs; edges represent GND ≤ 28%; node colors and titles represent 16S rRNA phylogenetic lineages. Text indicates GEN ID, 16S rRNA-based lineage, count of member SAGs and network linkage (ratio of actual versus possible link counts). Pane A contains all resolved GENs, pane B displays GEN #1 in greater detail. Note “*” indicates tentative identities of GENs that lack SAGs with 16S rRNA genes recovered.

**Fig. S5.**
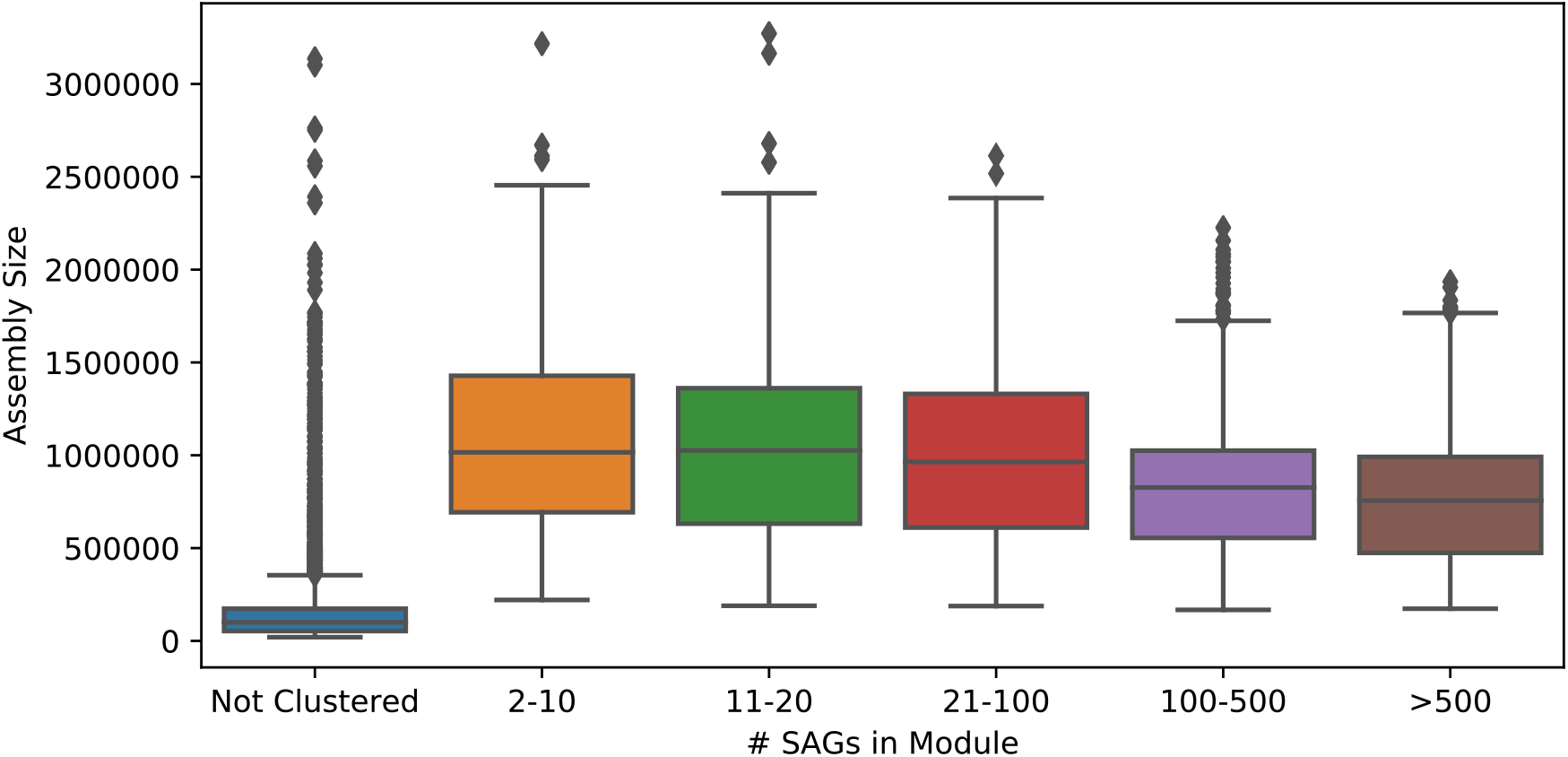
Genome assembly size distribution in GENs with diverse member count.

**Table S1. (separate file)**

Assignments of GORG-Tropics SAGs to GENs illustrated in Figs. 4 and S4.

**Table S2. (separate file)**

General characteristics of GENs illustrated in Figs. 4 and S4. Note “*” indicates tentative identities of GENs that lack SAGs with 16S rRNA genes recovered.

